# TREM2 mediates MHCII-associated CD4^+^ T cell response against gliomas

**DOI:** 10.1101/2023.04.05.535697

**Authors:** Jiaying Zheng, Lingxiao Wang, Shunyi Zhao, Wenjing Zhang, Yuzhou Chang, Aastha Dheer, Shan Gao, Shengze Xu, Katayoun Ayasoufi, Rawan Al-kharboosh, Manling Xie, Aaron J. Johnson, Haidong Dong, Alfredo Quiñones-Hinojosa, Long-Jun Wu

**Author notes:** Correspondence: Dr. Long-Jun Wu; Department of Neurology; Mayo Clinic; 200 First Street SW; Rochester, MN 55905.

## Abstract

Triggering receptor expressed on myeloid cells 2 (TREM2) was recently highlighted as a novel immune suppressive marker in peripheral tumors. The aim of this study was to characterize *TREM2* expression in gliomas and investigate its contribution in glioma progression by using *Trem2^-/-^*mouse line. Our results showed that higher *TREM2* expression was correlated with poor prognosis in glioma patients. Unexpectedly, TREM2 deficiency did not have a beneficial effect in a pre-clinical model of glioma. The increased *TREM2* expression in glioma was likely due to increased myeloid cell infiltration, as evidenced by our single-cell analysis showing that almost all microglia and macrophages in gliomas were TREM2^+^. Furthermore, we found that deficiency of TREM2 impaired tumor-myeloid phagocytosis and MHCII presentation, and significantly reduced CD4^+^ T cells in tumor hemispheres. Our results revealed a previously unrecognized protective role of tumor-myeloid TREM2 in promoting MHCII-associated CD4^+^ T cell response against gliomas.

**SUMMARY:** Authors found that although higher *TREM2* expression is correlated with poor prognosis in glioma patients, its absence has no beneficial effect in a pre-clinical model of glioma. Deficiency of TREM2 impairs myeloid cell phagocytosis of tumor debris, leading to a reduction in MHCII-dependent CD4^+^ anti-glioma immunity.

## INTRODUCTION

Antitumor immunity requires the presence of both major histocompatibility complex (MHC) class I and MHC class II (Alspach et al., 2019), which activate CD8^+^ and CD4^+^ cells, respectively, although their tumoricidal contributions vary across different types of tumors. For instance, in murine colon adenocarcinoma and sarcomas, anti-tumor immunity relies on CD8^+^ T cell infiltration and effectively respond to anti-programmed cell death protein 1 (PD-1) immunotherapy (Binnewies et al., 2021; Molgora et al., 2020; Oh et al., 2018). In contrast, in brain tumors, recent studies have suggested that MHC class II-restricted CD4^+^ tumor-infiltrating lymphocytes (TILs) play a key role in regulating tumor clearance (Chen et al., 2022; Kilian et al., 2022). As a results, anti-cytotoxic T-lymphocyte-associated antigen 4 (CTLA-4), but not anti-PD-1, extended the survival of glioma mice in a CD4^+^ T cell-dependent manner (Chen et al., 2022).

MHC class II-restricted antigen presentation requires antigen presenting cells to efficiently engulf and degrade exogenous antigenic material (Unanue, 2002). As a cell surface receptor, TREM2 is expressed exclusively in microglia (Colonna and Wang, 2016) in the central nervous system (CNS), macrophages (Do et al., 2022) and dendritic cells (DCs) (Bouchon et al., 2001) (Ulland and Colonna, 2018). The deficiency of TREM2 impairs the ability of microglia to sense and degrade numerous antigenic materials, including but not limited to pathogens (N’Diaye et al., 2009), apoptotic neurons (Takahashi et al., 2005), β-amyloid (Wang et al., 2015; Wang et al., 2016; Zhao et al., 2018), TAR DNA binding protein 43 (TDP-43) (Xie et al., 2022a), myelin (Cantoni et al., 2015), and neuronal synapses (Scott-Hewitt et al., 2020). Early studies reported that upon ligation of TREM2, genes related to MHC class II rapidly upregulate (Bouchon et al., 2001). Additionally, tumor-myeloid cells in CD4^+^ mediated glioma regression upregulated genes involved in pathways related to phagocytosis, including *Trem2* (Chen et al., 2022). Increased phagocytosis of tumor cells by macrophages has been shown to prolong the survival of glioma-burden mice (Zhai et al., 2021).

Interestingly, TREM2 deficiency reduced the immunosuppression of tumor-associated myeloid cells, suggesting that TREM2 can be detrimental in mouse models of peripheral tumors (Katzenelenbogen et al., 2020; Molgora et al., 2020; Timperi et al., 2022, Binnewies et al., 2021; Zhang et al., 2022). To this end, we sought to investigate whether TREM2 is beneficial or detrimental in brain tumors. Addressing this question can further our understandings of complicated immune responses in brain tumors. Here we found that while the TREM2 expression is positively associated with human glioma prognosis. Surprisingly, TREM2 deficiency in a classic glioma murine model did not slow disease progression. We further revealed that loss of TREM2 dampened the phagocytosis and MHCII antigen presentation. Our study highlighted the importance of myeloid TREM2 in promoting MHCII-associated CD4^+^ T cell response in gliomas.

## MATERIALS AND METHODS

### Animals

The *Trem2^-/-^* line was kindly provided by Dr. Marco Colonna at the Washington University School of Medicine, St. Louis, and was bred at Mayo Clinic (Xie et al., 2022a). Wild type mouse line was purchased from Jackson Laboratory. All animals were housed under standard conditions (21 - 22 °C; 55% humidity) in individually ventilated cages, with a 12-h light/dark cycle and ad libitum access to food and water. Male and female mice aged between 8 to 14 weeks were used in the studies. All experimental procedures were approved by the Mayo Clinic’s Institutional Animal Care and Use Committee (IACUC).

### Tumor cell culture

Murine GL261 glioma inoculation of C57BL/6 mice is a well-established experimental model of human glioblastoma (Haddad et al., 2021). GL261 is a syngeneic mouse model of glioblastoma in C57BL/6 mice that does not require an immunodeficient host (Jacobs, Valdes et al. 2011). The GL261 cell line transduced with firefly luciferase (GL261-luc) for *in vivo* monitoring of tumor kinetics was kindly provided by the laboratory of Dr. Aaron J. Johnson (Mayo Clinic, Rochester, MN). The cell line GL261-luc transduced with mCherry (GL261-luc-mCherry) for *in vivo* phagocytosis study was generated by the laboratory of Dr. Alfredo Quiñones-Hinojosa (Mayo Clinic, Jacksonville, FL). Cells were grown in Dulbecco’s modified Eagle medium (DMEM) (Gibco, #11965092) with 10% fetal bovine serum (FBS) (Sigma-Aldrich, #F2442) and 1% Penicillin-Streptomycin (Gibco, # 15140122), in a 37 °C humidified incubator with 5% CO_2_. For tumor inoculation, cells were dissociated with TrypLE™ Express (Gibco, # 12605010) and resuspended in phosphate buffered saline (PBS) at a final concentration of 4-6 × 10^4^ cells per μL.

Murine MC38 cell line derived from C57BL6 murine colon adenocarcinoma (Corbett et al., 1975) was kindly provided by the laboratory of Dr. Haidong Dong (Mayo Clinic, Rochester, MN). Cells were grown in Roswell Park Memorial Institute Medium (RPMI) (Corning, #10-040-CV) with 10% FBS and 1% Penicillin-Streptomycin, in a 37 °C humidified incubator with 5% CO_2_. For tumor inoculation, cells were dissociated with TrypLE™ Express, washed and resuspended in PBS at a final concentration of 5 × 10^3^ cells per μL.

### Inoculation of GL261 gliomas

Under isoflurane anesthesia, a 0.5-cm longitudinal incision was made on the scalp, and a burr hole was drilled using a high-speed dental drill (ML: ±1.5; AP: +1.5). Using a stereotactic frame, the needle of a Hamilton syringe was then lowered 3.5 mm into the striatum and a total of 4-6 × 10^4^ GL261-luc cells in 1-2 μL were injected, as previously described (Ayasoufi et al., 2020). The wound was closed using 6-0 ETHILON® Nylon Suture (Ethicon, #1660G).

To assess tumor burden in GL261-luc-bearing mice, bioluminescence imaging was used as previously described (Ayasoufi et al., 2020). Mice were intraperitoneally injected with 200 μL of 15 mg/μL D-Luciferin in PBS (Goldbio, #LUCK-1G), and anaesthetized with 2% isoflurane during imaging. Mice were scanned using the IVIS Spectrum system (Xenogen Corp.) at Mayo Clinic, running Living Image software.

### Inoculation of MC38 tumors

MC38 cells were washed and resuspended in sterile PBS, and then injected subcutaneously into previously shaved flanks of the mice. A total of 5 × 10^5^ cells in 100 μL PBS were injected into the mammary fat pad. Mice were monitored daily, and tumors were measured twice per week using calipers. All mice were sacrificed on day 22.

### RNA sequencing analysis

Single-cell RNA sequencing files containing human newly diagnosed GBM samples and mouse GL261 glioma samples were downloaded from the GEO database with accession number GSE163120 (Pombo Antunes et al., 2021). Seurat package (v4.3.0) was used for downstream analysis (Hao et al., 2021). In brief, Seurat objects were created from the feature-barcode matrices and annotated by the metadata file provided by the original study. The expression matrices were normalized with a scale factor of 1e6 and scaled. Top 200 variable features were calculated and used for linear dimension reduction by the principal component analysis (PCA). Harmony package (v0.1.1) was used to integrate data from each biological individual (Korsunsky et al., 2019). First 20 Harmony dimensions were used to calculate neighbor cells, and cell clusters were called by the FindCluster function with a resolution of 0.5. Cell identity of each cluster was determined based on the marker genes provided in the original study. UMAP of cells from human newly-diagnosed GBM samples was projected by the DimPlot function. TREM2 expression was plotted by the RidgePlot function. Normalized TREM2 expression matrix was extracted from the Seurat object and examined. The ProjecTILs package was used to determine the identity and ratio of T cells in the human newly diagnosed GBM samples (Andreatta et al., 2021).

Single-cell RNA sequencing files containing MC38 glioma samples from wild type and *Trem2^-/-^* mice were also downloaded from GEO database with accession number GSE151710 (Molgora et al., 2020). Seurat objects were created from the feature-barcode matrices and annotated by the metadata file provided by the original study. Tumor associated macrophages were first subset out according to the annotations described in the original paper and then re-annotated with the more detailed macrophage annotation file provided by the original study. The expression matrices were then normalized with a scale factor of 1e6 and scaled. Top 2000 variable features were calculated and used for PCA. Harmony package (v0.1.1) was used to integrate data from each biological individual. UMAP projection was done using first 50 dimensions from Harmony. The expression heatmap of selected genes were plotted by the FeaturePlot function in Seurat.

### Spectrum flow cytometry

Mice were perfused with 40 mL of 1 × PBS through intracardiac administration. After perfusion, tumor hemispheres were processed using a previously published protocol for enriching brain infiltrating immune cells, using the Dounce Homogenizer followed by centrifugation on a 30% Percoll (Sigma, P1644-1L) gradient (Cumba Garcia et al., 2016).

Zombie NIR viability dye (1:1000, BioLegend, 77184) was used to label dead cells. Dissociated cells were further incubated with a with a combination of the following antibodies along with a Fc blocking antibody, rat anti-CD16/CD32 (1:100, BD Pharmingen, 553142): BUV395 anti-F4/80 (1:100, BD Pharmingen, T45-2342), BUV615 anti-NK1.1 (1:100, BD Pharmingen, 751111), BUV805 anti-ICOS (1:100, BD Pharmingen, 568039), BV421 anti-CD62L (1:100, BioLegend, 115537), BV510 anti-CD4 (1:50, BioLegend, 100449), BUV570 anti-CD44 (1:100, BioLegend, 103037), BV605 anti-CTLA4 (1:100 intracellular staining, BioLegend, 106323), BV650 anti-Ly6C (1:100, BioLegend, 128049), BV750 anti-MHCII (1:500, BioLegend, 747458), BV785 anti-CD8α (1:200, BioLegend, 100750), FITC anti-PD1 (1:100, BioLegend, 135213), Spark Blue 550 anti-Ly6G (1:500, BioLegend, 127663), PerCP anti-CD45 (1:100, BioLegend, 103130), PE-Cy5 anti-CD11b (1:1000, Tonbo, 55-0112-U100), PE-Fire 700 anti-CD3 (1:200, BioLegend, 100272), APC anti-Foxp3 (1:50 intracellular staining, eBioscience, 17-5773-82), Spark NIR 685 anti-CD69 (1:100, BioLegend, 103277).

Samples were assessed by a spectral flow cytometer (Cytek Aurora, Cytek Biosciences) equipped with SpectroFlo software (Cytek Biosciences). Acquired flow cytometry results were analyzed by FlowJo software (BD Life Sciences).

### *In vivo* two-photon imaging

Craniotomy were performed previously described (Liu et al., 2019) (Eyo et al., 2018). In brief, under isoflurane anesthesia (3% induction, 1.5-2% maintenance), a circular craniotomy (<5 mm diameter) was made over somatosensory cortex with the center at about −2.5 posterior and +2 lateral to bregma. A total of 1-2 × 10^3^ GL261-luc-mCherry cells in 0.3 μL were injected into cortex. A circular glass coverslip (4 mm diameter, Warner) was secured over the craniotomy using dental cement (Tetric EvoFlow). A four-point headbar (NeuroTar) was secured over the window using dental cement. 7-14 days after surgery, we can observe mCherry tumor in the center of the window.

### Formalin-fixed paraffin-embedded (FFPE), immunofluorescence staining and confocal imaging

FFPE was used to embed endpoint tumor brains, to reduce autofluorescence of CD4 staining in tumor core regions. After 24-48 hours of fixation, tissues were dissected, placed in embedding cassettes. Fixed tissues were then transferred to 70% ethanol and processed as follows:; 70% ethanol for 1 h, 85% ethanol for 1 h, 95% ethanol for 3 × 30 min, 100% ethanol 3 × 30 min, xylene 3 × 30 min. After xylene, tissues were embedded into paraffin at 60°C across four changes, 2 × 45 min, 2 × 60 min. Tissues were immersed in liquid throughout the process.

Brain sections (5 µm) were obtained by a Leica microtome. Paraffin was removed from samples by consecutive 2 × 10 min washes with xylene. Xylene was then removed with graded washes of ethanol to water (100% ethanol 3 × 3 min, 96% ethanol 2 × 3 min, 85% ethanol 1 × 3 min, 70% ethanol 1 × 3 min, ddH_2_O 20s). The samples were immediately proceeded to antigen retrieval using Tris EDTA buffer (pH = 9) in 70 °C for 60 min. After cooling down, we removed the antigen retrieval buffer and washed slides with PBS.

For immunostaining, the slides were blocked by 4% of bovine serum albumin (EMD Millipore Corp, 126615-25mL) in PBS with 0.2% Tween-20. Primary antibodies were stained overnight in 4 °C with rat-anti-CD4 (1:100, Invitrogen, 14-9766-82) and rabbit-anti-Iba1 (1:500, Abcam, Ab178847). Slides were incubated with secondary antibodies of goat-anti-rat 488 and goat-anti-rabbit 594 (1:500, Invitrogen, A11006, A11037) 1.5 hours in room temperature. Sections were washed and mounted with DAPI Fluoromount-G mounting medium (SouthernBiotech). Fluorescent images were obtained by a confocal microscope (LSM 980, Zeiss). Cell counting was manually quantified, and brightness and contrast were adjusted by ImageJ (National Institutes of Health).

### RNA extraction and quantitative **reverse** transcription polymerase chain reaction (qRT-PCR)

RNA was extracted from the endpoint hemispheres using RNAeasy Plus Mini Kit (Qiagen, 74134). Reverse transcription of RNA was performed suing iScript™ cDNA synthesis Kit (Bio-Rad, 1708891). cDNA was added to a reaction mix (20 μL final volume) containing gene-specific primers and SYBR Green Supermix (Bio-Rad, 1725271). All samples were run in duplicates in LightCycler 480 II (Roche). The relative gene expression was normalized to *Gapdh* and assessed using the 2^−ΔΔCT^ method. Primer sequences and information are as follows (5′ - 3′): *Gapdh*: CATCTTCCAGGAGCGAGACC (forward), TCTCGTGGTTCACACCCATC (reverse); *Trem2*: CTCCAGGAATCAAGAGACCTCC (forward), CCGGGTCCAGTGAGGATCT (reverse); *Cd68*: TGTCTGATCTTGCTAGGACCG (forward), GAGAGTAACGGCCTTTTTGTGA (reverse); *H2-Aa*: TCAGTCGCAGACGGTGTTTAT (forward), GGGGGCTGGAATCTCAGGT (reverse); *Cd4*: AGGTGATGGGACCTACCTCTC (forward), GGGGCCACCACTTGAACTAC (reverse).

### Statistical analysis

GraphPad Prism 9 was used for statistical analysis. All results were reported as mean with standard error of the mean (SEM). Differences between groups were measured by a two-tailed t test or two-way ANOVA. Mice were of mixed sexes. Mice within experiments were age and sex matched.

## RESULTS

### High *TREM2* expression correlates with poor prognosis in human brain tumors

To understand the potential relevance of TREM2 in cancer, first we explored the *TREM2* expression profile across multiple tumor samples through the GEPIA portal (Tang et al., 2017). Interestingly, we found a prevalent increase of *TREM2* in 22 tumor types compared to the paired normal tissues (**Supplementary Table 1**). Brain tumors exhibited remarkably elevated levels of *TREM2* expression compared to other tumor types (GBM: median = 117.85 TPM; LGG: median = 54.32 TPM) (**Figure 1A**). Although recent studies have demonstrated that *TREM2* is linked to worse outcomes in peripheral murine tumors, including sarcoma, colon adenocarcinoma and breast adenocarcinoma (Katzenelenbogen et al., 2020; Molgora et al., 2020; Timperi et al., 2022), the role of increased *TREM2* expression in brain tumors has not been elucidated. Therefore, we next explored the correlation between *TREM2* expression and brain tumor prognosis using The Cancer Genome Atlas (TCGA) database, GlioVis data portal (Bowman et al., 2017). Analysis showed that *TREM2* expression is increased with the severity of glioma WHO grades, ranging from Grade II to IV (**Figure 1B**). To assess the clinical relevance of *TREM2* expression and its prognostic potential, we explored its association with clinically recognized molecular classifications of glioma. It is known that mutations in isocitrate dehydrogenase (IDH) (a mutation in either *IDH1* or *IDH2*) and deletion of chromosome arms 1p and 19q (1p/19q) are strong prognostic biomarkers associated with improved survival in gliomas (Cancer Genome Atlas Research et al., 2015; Zhao et al., 2014). Accordingly, our analysis revealed relatively lower *TREM2* expression in the IDH mutant and 1p/19q codeleted cases when compared to IDH wild-type and 1p/19q non-codeleted cases, respectively (**Figure 1C & 1D**). We further evaluated the *TREM2* implication on overall survival outcomes using the TCGA and Chinese Glioma Genome Atlas (CGGA) databases. With a threshold of 75% quantile, high *TREM2* expression correlated with worse overall survival in both glioma cohorts (**Figure 1E & 1F**). Collectively, these data suggested that increased *TREM2* expression is strongly associated with poor prognosis in brain tumors.

**Figure 1:**
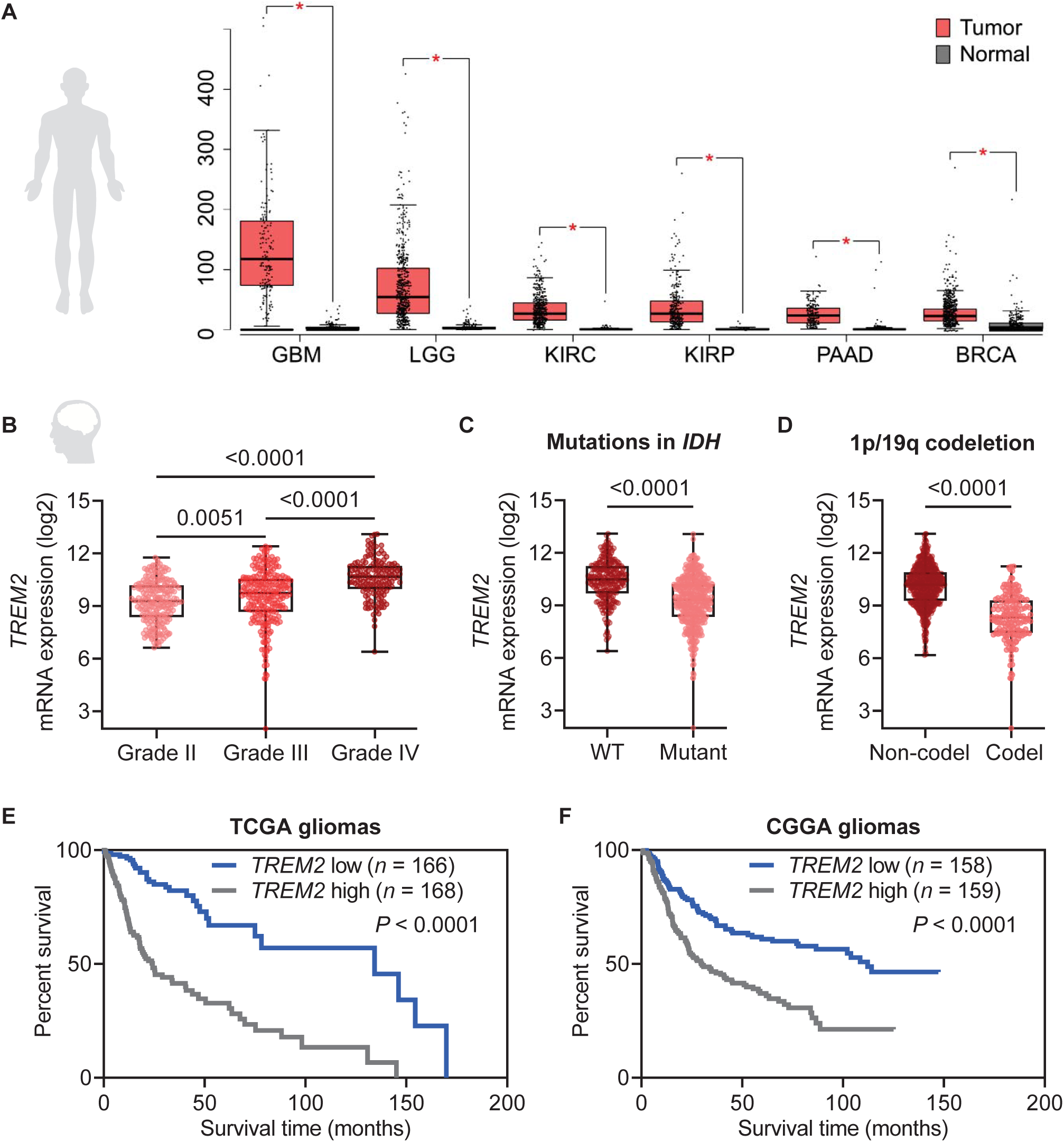
High *TREM2* mRNA expression in human gliomas is associated with poor patient prognosis. **A.** Top six types of tumors with elevated *TREM2* expression in a descending order from left to right, which were glioblastoma (GBM), brain lower grade glioma (LGG), kidney renal clear cell carcinoma (KIRC), kidney renal papillary cell carcinoma (KIRP), pancreatic adenocarcinoma (PAAD), breast cancer (BRCA). **B.** Gene expression of *TREM2* obtained from TCGA_GBMLGG dataset (*n* = 620). *TREM2* expression was increased along with glioma WHO grade (grade II, *n* = 226, median = 9.28; grade III, *n* = 244, median = 9.74; grade IV, *n* = 150, median = 10.67). **C.** *TREM2* expression was relatively lower in the IDH mutant (mutant, *n* = 429, median = 9.32; WT, *n* = 233, median = 10.49). **D.** *TREM2* expression was relatively lower in the chromosome 1p/19q codeletion group (codel, *n* = 169, median = 8.33; non-codel, *n* = 494, median = 10.20). **E & F.** Kaplan-Meier survival curves generated for *TREM2* expression in glioma patients. Patients were divided in high- and low-expressing groups based on quantile of *TREM2* expression. In the TCGA dataset, higher *TREM2* expression correlated with worse overall (*TREM2* high, *n* = 168, events = 92; median = 24.2; *TREM2* low, *n* = 166, events = 31; median = 134.3). Similar result was observed from the CGGA dataset (*TREM2* high, *n* = 159, events = 103; median = 30.1; *TREM2* low, *n* = 158, events = 66; median = 112.1). Data were tested for normal distribution using Shapiro-Wilk test first. *P*-values were acquired using two-tailed Student t-tests if data were normally distributed, or Mann-Whitney test if not. Survival curves were analyzed using log-rank test.

### TREM2 deficiency accelerates tumor progression in brains but not periphery

Given the strong association between TREM2 and glioma prognosis in patients, we investigated whether TREM2 contributes to glioma progression. To this end, we used an immunocompetent glioma model by inoculating GL261 cells into the brains of both WT and *Trem2^-/-^* mice (**Figure 2A**). At humane endpoints, there was no significant difference in tumor size between WT and *Trem2^-/-^*mice, with the weight of tumor hemispheres being at least twice that of contralateral hemispheres (**Figure 2B**). We then examined TREM2 expression in both tumor hemispheres and contralateral hemispheres of glioma endpoint mice. Our results showed a significant increase of *Trem2* expression in WT tumor hemispheres compared to contralateral hemispheres. Additionally, *Trem2* was not detected in either tumor or contralateral hemispheres of *Trem2^-/-^*mice, demonstrating the absence of *Trem2* expression in GL261 cells (**Figure 2C**).

**Figure 2:**
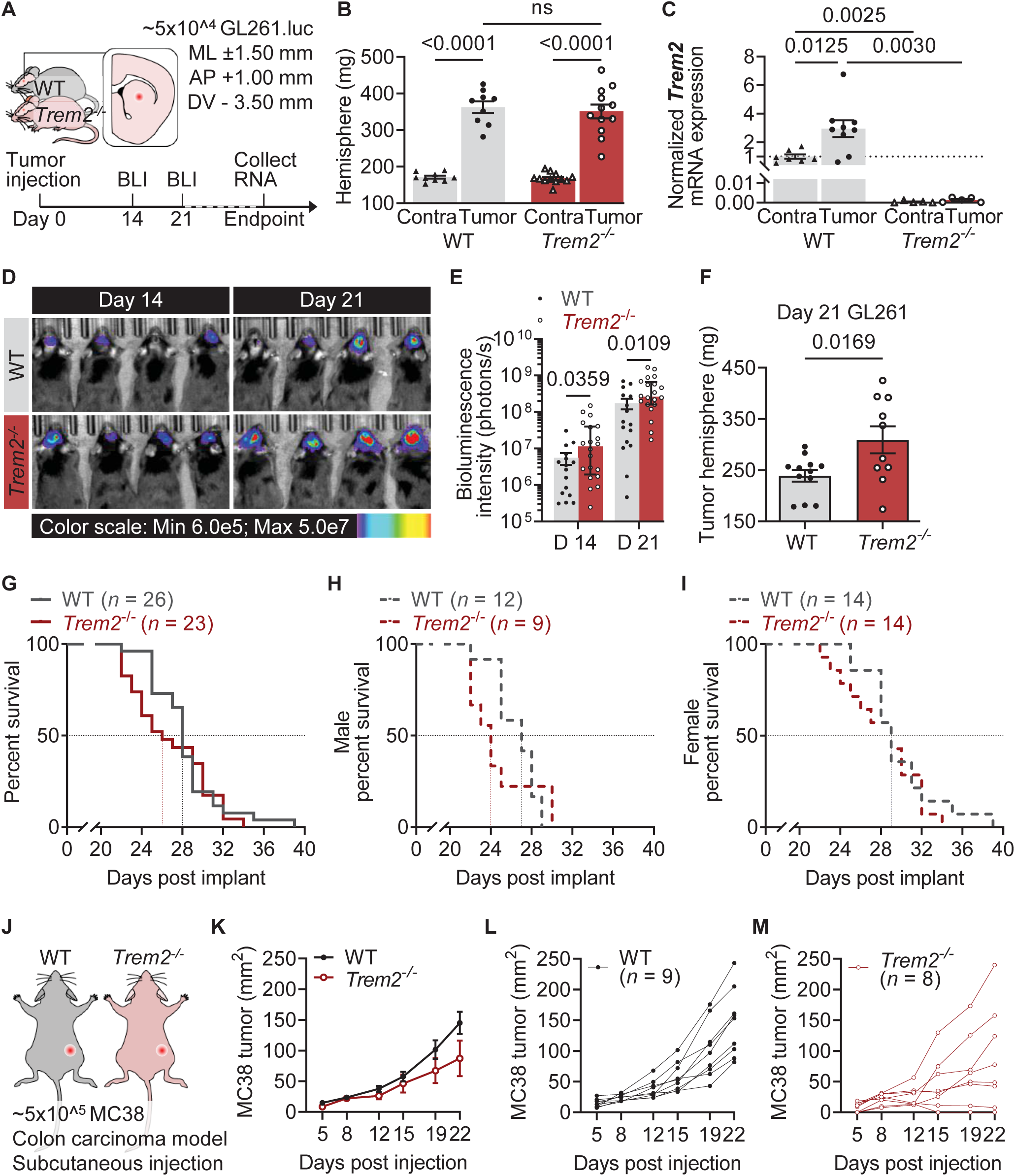
TREM2 deficiency accelerates glioma but not peripheral tumor progression. **A.** A schematic illustration of establishing an immunocompetent glioma model using murine glioma GL261 cells. Tumor size was monitored by bioluminescence imaging (BLI) every 7 days from day 14 post-inoculation. When mice reached the humane endpoint, contralateral and tumor hemispheres were collected separately for further analysis. **B.** The weight of hemispheres of WT and *Trem2^-/-^*when humane endpoints were reached. **C.** The mRNA levels of *Trem2* in the contralateral and tumor hemispheres were quantified by qRT-PCR. **D & E.** Representative bioluminescence images and statistical analysis showed that brain tumor burden was relatively higher in *Trem2^-/-^* mice compared to WT mice. **F.** A larger tumor size was observed in *Trem2^-/-^*mice compared to the WT mice 21 days after tumor inoculation, as indicated by the increased weight of tumor hemisphere. **G - I.** The survival study using 26 WT (12 males and 14 females) and 23 *Trem2^-/-^* (9 males and 14 females) mice showed Trem2 deficiency did not confer any survival benefit in glioma. **J - M.** MC38 subcutaneous tumor experiment using 9 WT and 8 *Trem2^-/-^* mice showed a clear trend (*P* = 0.0575) towards an attenuated tumor progression in *Trem2^-/-^* mice. The data were shown as mean ± SEM. The data were tested for normal distribution using Shapiro-Wilk test first. *P*-values were acquired using two-tailed Student *t*-tests or two-way ANOVA if the data were normally distributed, or Mann-Whitney test if they were not. Survival curves were analyzed using log-rank test.

Unexpectedly, bioluminescence imaging revealed a higher burden of brain tumors in *Trem2^-/-^* mice compared to WT animals at both day 14 and day 21 post-tumor (**Figure 2D & 2E**). Consistent with the bioluminescence result, a larger tumor size was observed in *Trem2^-/-^* mice compared to the WT mice, as indicated by the increased weight of tumor hemisphere at day 21 post-tumor inoculation (**Figure 2F**). A survival study using 26 WT (14 females, 16 males) and 23 *Trem2^-/-^*mice (14 females, 9 males) showed that Trem2 deficiency did not benefit glioma survival, and even led to a slightly shorter medium survival time compared to the WT (**Figure 2G**). This conclusion was consistent regardless of whether the data was analyzed by sex. (**Figure 2H & 2I**). This surprising phenotype is opposite to that observed in multiple peripheral tumor models showing slower tumor progression in *Trem2^-/-^* mice (Katzenelenbogen et al., 2020; Molgora et al., 2020; Timperi et al., 2022). To discern whether the discrepancy was potentially due to the tumor types, we repeated the same experiments using MC38 subcutaneous model with 9 WT and 8 *Trem2^-/-^*mice (**Figure 2J**). Similar to previous findings (Molgora et al., 2020), we observed a clear trend towards attenuated tumor progression in *Trem2^-/-^* mice (**Figure 2K**). Individual plots showed variations in *Trem2^-/-^* group, and the attenuated trend is contributed by *Trem2^-/-^* mice having tumor regression (**Figure 2L & 2M**). Taken together, these results indicate that TREM2 may have unrecognized protective roles specific to brain tumors, highlighting the need for further investigation.

### TREM2 is highly expressed in tumor-associated microglia and macrophages

To investigate the potential protective role of TREM2 in glioma, we first queried TREM2 expression patterns at the cellular level using recently published glioma dataset (Pombo Antunes et al., 2021). When tumor occurs, there is a massive infiltration of immune cells into the brain. We examined a total of 21,303 cells from human newly diagnosed glioblastoma (GBM) and 27,276 WT cells from mouse GL261 gliomas at day 21 post inoculation to map *TREM2* transcription by different cell populations. Tumor-associated macrophages (TAMs) were found to be the largest immune cell population in both newly diagnosed GBM and GL261 glioma, comprising approximately 80% and 50% of the immune cells, respectively (**Figure 3A & 3C**). TAMs were composed of two main populations, microglia and macrophages. Notably, in newly diagnosed GBM, microglia accounted for a larger proportion (58.51%) than macrophages (23.35%) (**Figure 3A**), whereas in mouse glioma, the proportion of macrophages was much higher (49.98%) compared to microglia (7.85%) %) (**Figure 3C**). Among all immune cell populations, *TREM2* was expressed in almost all microglia, with the highest levels of transcription observed in these cells. TREM2 expression was also detected in multiple subtypes of macrophages (**Figure 3B & 3D**). Additionally, TREM2 expression was found to be present in 66.49% of human DCs, and to a lesser extent in other immune cell populations or mouse DCs (28.34%).

**Figure 3:**
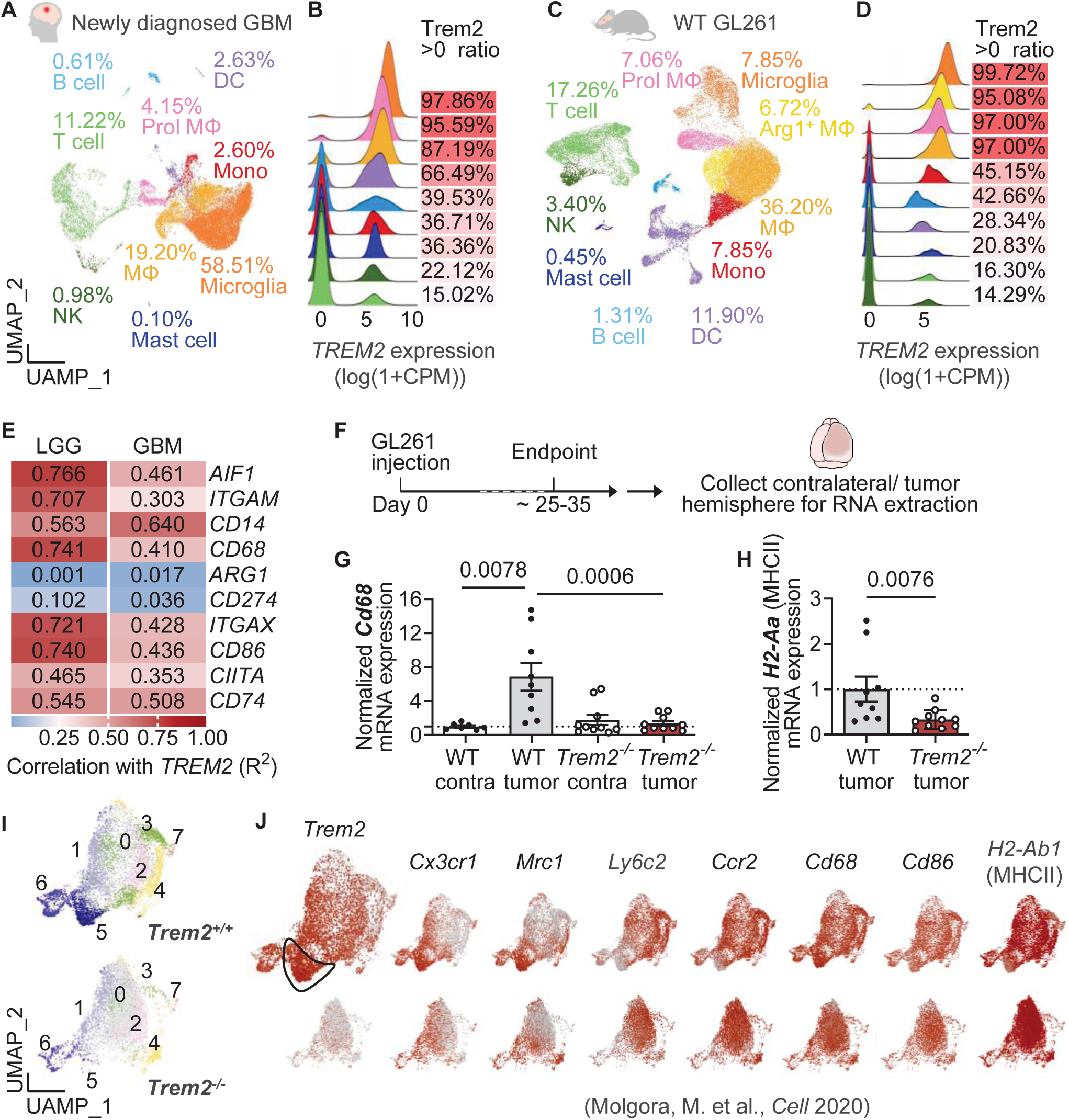
TREM2 deficiency dampens MHC class II expression. **A & C.** UMAP plots displaying the immune cells in patients with GBM and in mice with glioma GL261. **B & D.** *TREM2* transcription in different immune cell populations. **E.** The correlation between *TREM2* expression and signature genes related to myeloid cell phagocytosis, immunosuppression *t*, and antigen presentation was evaluated in LGG and GBM patients. The Spearman’s correlation test produces both *P*-values and correlation coefficients (R^2^). All listed genes had a *P*-value less than 0.05. Genes with an R^2^ value greater than 0.25 were considered to have a correlation (either positive or negative) with *TREM2* expression. **F.** Diagram of investigating the role of TREM2 in regulation of phagocytosis and antigen presentation in the mouse GL261 model. **G & H.** The mRNA levels of *Cd68* and *H2Aa* (encoding MHC class II) in the contralateral and tumor hemispheres were quantified by qRT-PCR. **I.** UMAP plots of myeloid clusters in the MCA tumors from *Trem2^+/+^* and *Trem2^-/-^*male mice. **J.** Feature plots of selected cluster markers and feature genes involved in phagocytosis and antigen presenting. The data were shown as mean ± SEM. The data were tested for normal distribution using Shapiro-Wilk test first. *P*-values were acquired using two-tailed Student *t*-tests if the data were normally distributed, or Mann-Whitney test if they were not.

We moved to TCGA dataset with a larger patient cohort to evaluate the correlation of *TREM2* expression with phagocytosis and antigen presentation features. Our analysis of human gliomas (LGG and GBM) revealed a strong correlation between *TREM2* expression level and tumor-associated myeloid markers, such as *AIF1* (encoding IBA1), *ITGAM* (encoding CD11B), and *CD14* (**Figure 3E**). This indicated that more myeloid infiltration resulted in higher *TREM2*. *CD68*, encodes a heavily glycosylated glycoprotein predominantly expressed in late endosomes and lysosomes of macrophages, was also strongly correlated with *TREM2*. Interestingly, *Arg1* (encodes arginase), and *CD274* (encodes PD-L1, an immune inhibitory receptor ligand) which are markers of immunosuppressive features, were poorly associated with *TREM2*. Genes involved in antigen presenting pathway such as *ITGAX* (encodes integrin alpha X chain protein CD11C, a marker of antigen presenting cells), *CD86* (provides costimulatory signals necessary for T cell activation and survival), *CIITA* (MHC class II transactivator, responsible for turning on MHC class II gene transcription), and *CD74* (the invariant chain required for the proper folding and trafficking of MHC class II in antigen presenting cells), were well-correlated with *TREM2* expression.

### TREM2 deficiency causes a partial loss of myeloid cells with phagocytic and antigen-presenting features

The observed correlation between *TREM2* expression and phagocytic and antigen-presenting markers in human gliomas prompted an *in vivo* exploration to assess the extent of TREM2’s impact on promoting a glioma-mediated immune response. We tested this idea using our GL261 mouse models in WT and *Trem2^-/-^* mice. We extracted RNA from tumor hemispheres and contralateral hemispheres at the endpoint of the mouse survival study when the glioma reached its maximum size (**Figure 3F**). We consistently found that *Cd68* expression in the tumor hemispheres was significantly higher than that in the contralateral hemispheres in WT mice, whereas TREM2 deficiency suppressed the elevation of *Cd68* expression in the tumor hemispheres (**Figure 3G**). The H2Aa, which encodes mouse class II antigen A, was also significantly lower in the *Trem2^-/-^* tumor hemispheres than the WT ones (**Figure 3H**).

We further investigated how TREM2 impacts the tumor-myeloid antigen presentation at the cellular level. We clustered WT and *Trem2^-/-^*tumor-associated myeloid cells from day 10 MCA/1956 tumors into 8 subsets using Uniform Manifold Approximation and Projection (UMAP) (**Figure 3I**). We found that Cluster 5 (CX3CR1-Macs) and Cluster 6 (Cycling-Macs), which were poorly represented in *Trem2^-/-^* mice, exhibited particularly high TREM2 expression. These two clusters showed a more resident macrophage-like profile with high levels of *Cx3cr1* and *Mrc1*, but low levels of *Ly6c2* and *Ccr2*. They also expressed lysosome markers *Cd68*, and antigen presenting cell markers *Cd86* and *H2-Ab1* (**Figure 3J**). This indicated TREM2 deficiency may lead to the loss of partial myeloid cells with antigen presenting features. Such antigen presenting myeloid cells may be critical to anti-glioma immunity.

TREM2 is a crucial player for phagocytosis function of myeloid cells (Colonna and Wang, 2016). To directly determine whether TREM2 deficiency impairs myeloid cell phagocytosis in brain tumor, we took advantage of *in vivo* two-photon imaging approaches. After transducing the GL261 tumor cell line with mCherry^+^ (red) labeling, we injected them into the somatosensory cortex of *Cx3cr1^Gfp/+^* mice after craniotomy (**Figure 4A**). Between 7-14 days after tumor inoculation, we observed CX3CR1^GFP^ myeloid cells closely interacting with tumor cells in the tumor core region. Interestingly, some of these myeloid cells were observed to uptake mCherry^+^ tumor debris and persisted for hours (**Figure 4B**). This mCherry^+^ signals can also be detected by flow cytometry (**Figure 4C**). We found the mCherry signals were colocalized with MHCII^+^F4/80^+^ antigen presenting like macrophages, with a higher percentage in WT compared to *Trem2^-/-^*mice ((**Figure 4D & 4E**). To examine the impact of TREM2 deficiency on the immune components of glioma, we performed flow cytometry on day 25 when most mice had a high tumor burden. Unbiased UMAP clustering of high parameter flow cytometry data obtained from total immune cells (CD45^+^) of perfused tumor hemispheres identified seven main clusters, including microglia, infiltrating myeloid, CD8 ^+^ T cells, natural killer cells (NK), B cells, regulatory T (Treg), and conventional T helper (Th) cells (**Figure 4F**). As the tumor size increased, the percentage of microglia in WT mice decreased (**Figure 4G**). However, the percentage of microglia was already low in *Trem2^-/-^* mice and further decreased with increasing tumor size (**Figure 4G**). This observation suggests that *Trem2^-/-^* microglia may have impaired activation in response to the tumor at the onset, leading to a lower proportion of microglia in the tumor microenvironment. The higher percentage of infiltrating cells in *Trem2^-/-^* mice may be a compensatory response to the impaired microglia activation. In infiltrating immune cells (CD45^hi^), infiltrating myeloid, NK cells and lymphocytes weighted similarly in WT and *Trem2^-/-^*(**Supplementary Figure 1A-1D**). WT and *Trem2^-/-^* showed a similar trend of increased percentage of infiltrating myeloid, and a decreased percentage of NK and lymphocytes along with increased tumor size. We further delved into the influence of TREM2 on infiltrating myeloid subsets. Consistent with findings of TREM2 deficient macrophages in MCA/1956 tumors (**Figure 3J**), a Ly6C^neg^F4/80^+^ cluster with high CD68 and MHCII expression was reduced in *Trem2^-/-^*mice (**Figure 4H**). This MHCII^hi^ macrophage-like subset is positively correlated with the tumor size. However, the percentage of this antigen-presenting-like macrophage subset was much higher in WT, particularly in high tumor burden hemispheres (more than 250 mg), compared to that in *Trem2^-/-^* mice (**Figure 4I-4J**). These results indicate that TREM2 deficiency reduces tumor-associated myeloid cells, particularly those antigen-presenting macrophage subsets.

**Figure 4:**
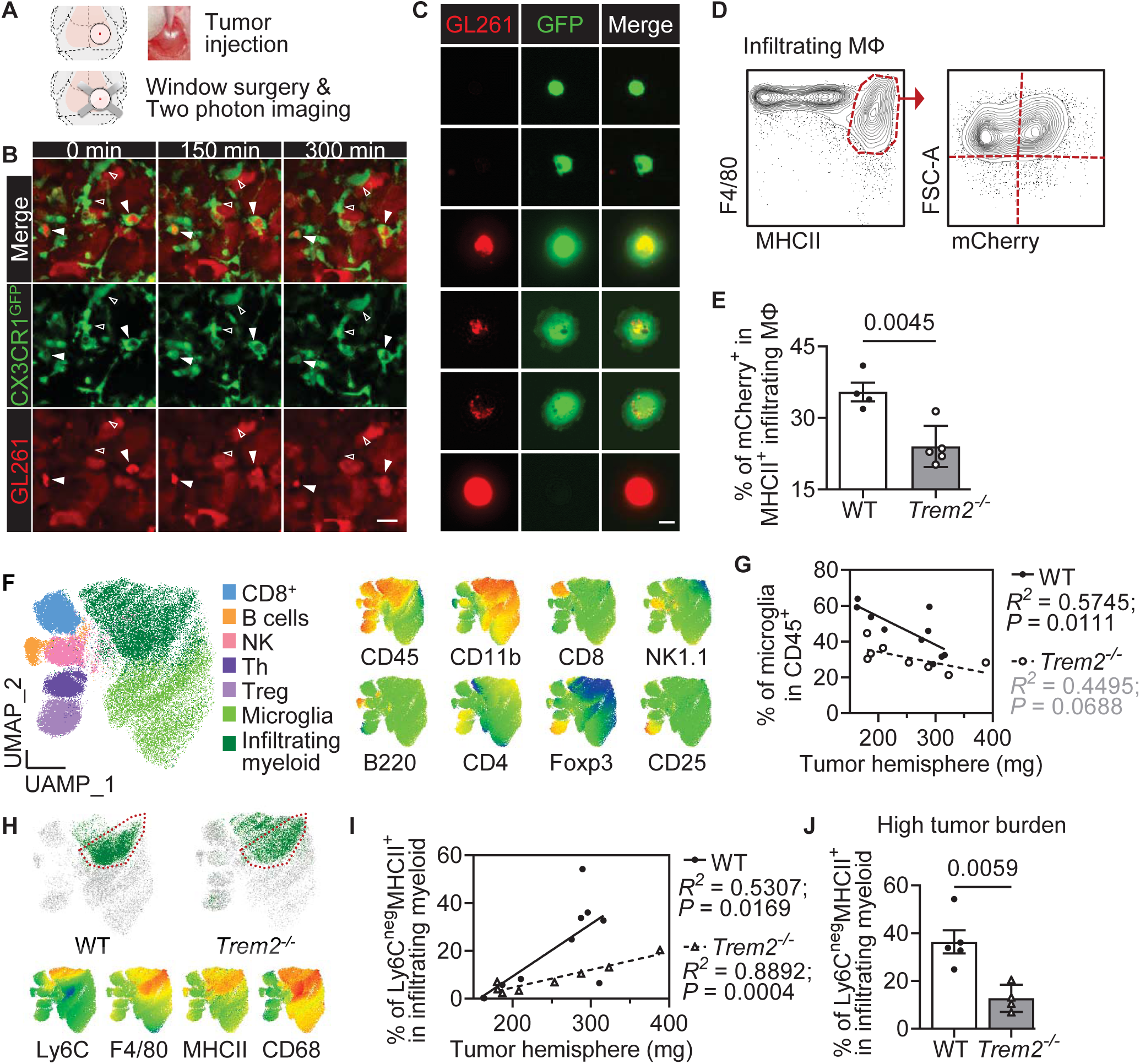
TREM2 deficiency impairs myeloid cell uptake tumor debris and antigen presentation. **A.** A schematic illustration of tumor inoculation and window surgery for *in vivo* two-photon imaging. **B.** *In vivo* imaging of CX3CR1^GFP^ (green) myeloid cells interacting with mCherry^+^ (red) tumor cells. Solid triangles indicated myeloid cells that uptake red tumor debris and hollow triangles indicating those that do not. Scale bar: 10 µm. **C.** Dissociated CX3CR1 ^GFP^ cells from brain tumor under EVOS microscope, with some containing red tumor debris. Scale bar: 10 µm. **D.** Flow cytometry gating showing the detection of mCherry^+^ tumor debris signal in F4/80^+^MHCII^+^ macrophages. **E.** The percentage of tumor debris signal in F4/80^+^MHCII^+^ macrophages. **F.** UMAP plots of CD45^+^ immune populations detected in the GL261 tumor hemispheres and selected cluster markers. **G.** Correlation between percentage of microglia in CD45^+^ and the weight of tumor hemispheres in WT and *Trem2^-/-^*. **H.** UMAP plots of infiltrating myeloid clusters in WT and *Trem2^-/-^* and selected myeloid cell markers. **I.** Correlation between percentage of Ly6C^neg^F4/80^+^ in infiltrating myeloid cells and the weight of tumor hemispheres in WT and *Trem2^-/-^*. **J.** Percentage of Ly6C^neg^F4/80^+^ in infiltrating myeloid cells in the high tumor burden group (tumor hemisphere > 250 mg). The bar graphs were shown as mean ± SEM. The data were tested for normal distribution using Shapiro-Wilk test first. *P*-values were acquired using two-tailed Student *t*-tests if the data were normally distributed, or Mann-Whitney test if they were not. For the correlation study, the Spearman’s correlation test produces both *P*-values and correlation coefficients (R²). A correlation is considered significant when the R² is greater than 0.25 and the *P*-value is less than 0.05.

### TREM2 deficiency impairs CD4^+^ T cell responses to gliomas

Considering CD4^+^ T cells’ engagement with antigen-presenting cells, we next determine whether TREM2 deficiency in the mouse model of glioma could impact CD4^+^ T cell infiltration. Indeed, we found that *Cd4* expression in WT glioma-endpoint hemispheres was higher compared to that in *Trem2^-/-^*mice (**Figure 5A**). This was corroborated by immunofluorescence staining of CD4 at the endpoint hemispheres, which consistently demonstrated a greater number of CD4^+^ TILs in WT mice than in *Trem2^-/-^* mice (**Figure 5B**). In addition, we found that CD4^+^ T cells mostly located in the tumor core region and were in contact with macrophages (Iba1^+^, ramified structure) with varying proximities and interactions (**Figure 5C**). In WT, approximately 20% of CD4^+^ TILs had no cell to cell contact with macrophages, 20% had some interaction via the tips of macrophage processes, 60% had tight interaction between cell soma, and 10% were enclosed by multiple macrophages. In *Trem2^-/-^* mice, CD4^+^ TILs had decreased intermediate contact with macrophages and increased enclosed type interaction (**Figure 5C-5D**). The formation of intermediate/tight contacts between T cells and antigen-presenting cells is thought to be a result of antigen stimulation (Stock et al., 2019), whereas the enclosed structure is likely to represent macrophages uptake of exhausted T cells. Thus, these results suggest that TREM2 in the antigen-presenting like macrophage subset is critical for the interaction with CD4^+^ TILs in gliomas.

**Figure 5:**
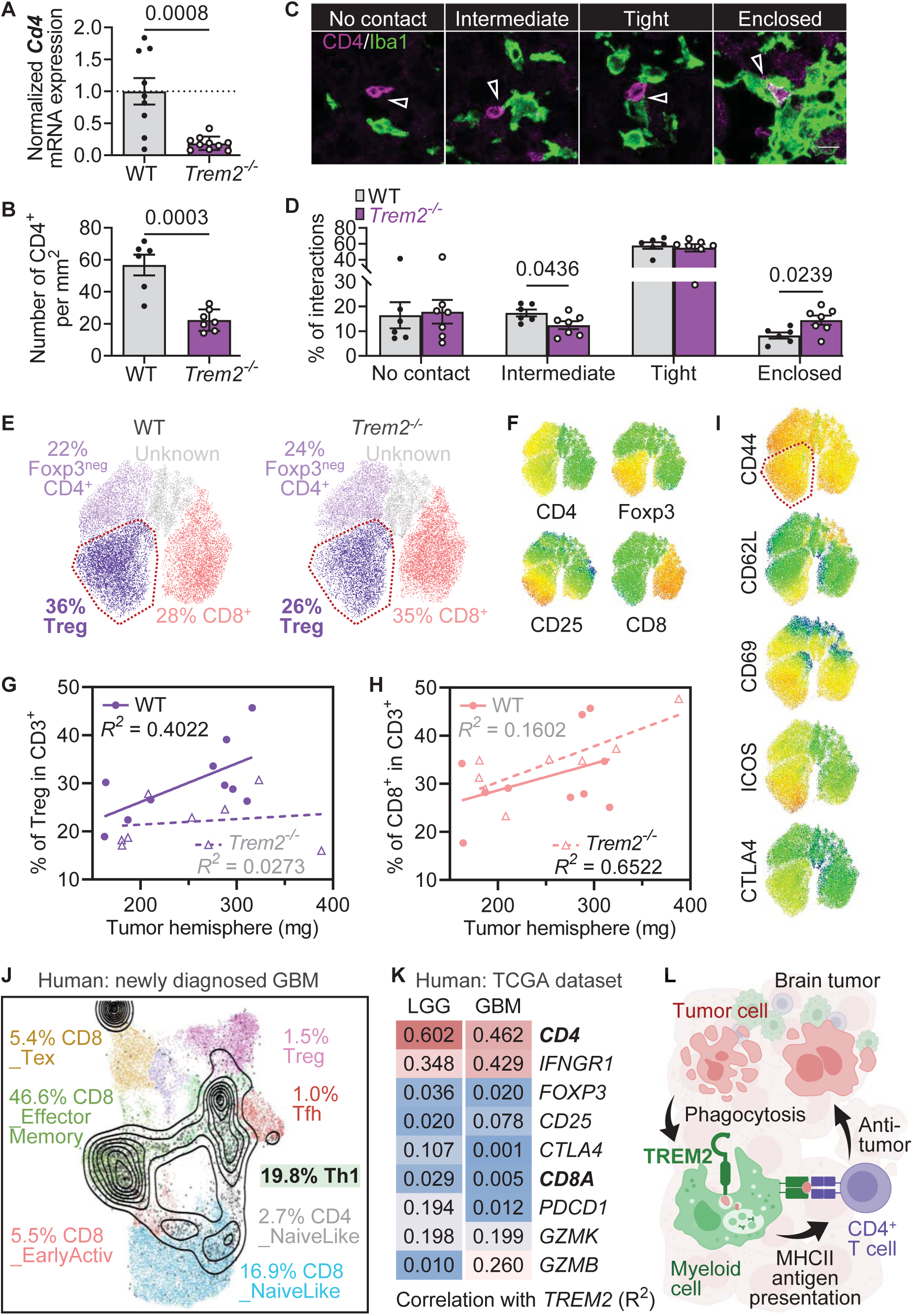
TREM2 is necessary for accumulation of CD4^+^ T cells in brain tumors. **A.** Quantification of *CD4* mRNA levels in tumor hemispheres using qRT-PCR. **B.** Quantification of the number of CD4^+^ T cells per mm^2^ in the tumor core using confocal microscopy on 5 µm thick brain slides. **C.** Representative images of CD4^+^ T cell and myeloid cell (Iba1^+^) interactions. Scale bar: 10 µm. **D.** Quantifications of different types of CD4^+^-myeloid cell interactions in WT and *Trem2^-/-^*. **E.** UMAP plots of CD3^+^ T cells in WT and *Trem2^-/-^*. **F & I.** Expression levels of T cell cluster markers and feature genes indicating T cell activation or immunosuppression. **G & H.** Correlation between percentage of T regulatory cell and CD8^+^ in total T cells (CD3^+^) and the weight of tumor hemispheres in WT and *Trem2^-/-^*. Spearman’s correlation test was used to calculate *P*-values and correlation coefficients (R^²^). **J.** Projection of T cells from newly diagnosed GBM using the ProjecTIL package to reveal T cell subsets. **K.** Correlation between *TREM2* expression and marker genes of T cell subtypes in LGG and GBM patients. All listed genes, except for *CTLA4*, had a *P*-value less than 0.05. **L.** Our working model proposes that TREM2-mediated phagocytosis of glioma debris by myeloid cells leads to further MHC class II presentation to CD4^+^ T cells, ultimately contributing to anti-tumor immunity in brain tumors. The bar graphs were shown as mean ± SEM. The data were tested for normal distribution using Shapiro-Wilk test first. *P*-values were acquired using two-tailed Student *t*-tests if the data were normally distributed, or Mann-Whitney test if they were not.

Surprisingly, upon analyzing TIL composition using flow cytometry, we found that the higher number of CD4^+^ TILs in WT mice was primarily due to Treg (CD25^+^Foxp3^+^). Only in WT mice was the increased proportion of Treg in total T cells (CD3^+^) correlated with the increased tumor size (**Figure 5E-5G**). The proportion of Treg in total T cells was also higher in WT mice than in *Trem2^-/-^*mice. There was no significant difference in the proportion of CD8^+^ TILs between WT and *Trem2^-/-^*, although the proportion of CD8^+^ TILs was positively corelated with tumor size in *Trem2^-/-^* mice (**Figure 5H**). We further investigated various T cell markers and found that the Treg cluster expressed a mixture of markers that positively regulated T cell activation (Beltra et al., 2020; Duhen et al., 2022) such as CD44^+^, ICOS^+^, CD69^+^, CD62L^+^, as well as negative regulatory markers such as CTLA4^+^ (**Figure 5I**). Therefore, this population could potentially have both anti- and pro-tumoral functions.

To further address the significance of CD4^+^ TILs in human gliomas, we analyzed T cell composition in newly diagnosed GBM. We projected total GBM T cells (Pombo Antunes et al., 2021) via ProjecTIL package (Andreatta et al., 2021) to reveal T cell subsets. We found that Th1-like CD4^+^ TILs were the second largest population in the newly diagnosed GBM T cells after effector memory CD8^+^ TILs, accounting for 19.8% of the total GBM T cells (**Figure 5J**). Furthermore, analysis of TCGA dataset revealed a correlation between *TREM2* and Th1-like markers such as *CD4* and IFN-gamma receptor 1 (*Ifngr1*) in both LGG and GBM (**Figure 5K**). However, no significant correlation was found between TREM2 expression and Treg markers such as *FOXP3* and *CD25* (also known as *IL2RA*), or immunosuppressive markers such as *CTLA4*. Additionally, markers of effector memory CD8^+^ TILs such as *PDCD1* and granzymes (mostly *GZMK*, with an intermediate level of *GZMB*) showed minimal correlation with TREM2 expression. Collectively, these results suggest a positive role of TREM2 in mediating MHCII-restricted CD4^+^ responses to gliomas. Our findings may have important clinical implications for the development of novel immunotherapeutic strategies targeting TREM2 and other myeloid-specific proteins in cancer treatment.

## DISCUSSION

In this study, we showed that while TREM2 was strongly corelated with poor prognosis in brain tumor patients, constitutional depletion of TREM2 did not result in beneficial effect, as demonstrated by our pre-clinical model of glioblastoma. Although there are species differences, our results indicate that increased TREM2 expression in glioma may not be the causal driver of tumor progression. Our single-cell analysis reveals that almost all microglia and macrophages in the GBM expressed TREM2, suggesting that increased *TREM2* expression may be a result of increased myeloid cell infiltration. It is the higher proportion of myeloid infiltration that results in a more immunosuppressive microenvironment, aggravating glioma progression (Zhang et al., 2019), rather than TREM2.

Our results further implied that there are differences in tumor immunity between the brain and peripheral tissues. In pre-clinical models of colon carcinoma and melanoma, CD8^+^ T cells have been shown to be the primary mediators of tumor reduction, and their depletion has been found to eliminate the protective benefits of both genetic and immunotherapeutic interventions (Ji et al., 2021; Katkeviciute et al., 2021). Additionally, in metastatic melanoma patients who responded to PD-1 blockade treatment, tumor regression was accompanied by the proliferation of CD8^+^ TILs (Tumeh et al., 2014). In this paradigm, TREM2 deficiency reduced the immunosuppressive activity of myeloid cells, which in turn led to improved preservation and functionality of CD8^+^ T cells responding to anti-PD-1; as a result, overall survival in mice was improved in *Trem2^-/-^* compared with WT mice (Binnewies et al., 2021; Katzenelenbogen et al., 2020; Molgora et al., 2020; Timperi et al., 2022). However, anti-PD-1/PD-L1 immunotherapy has shown limited efficacy in most clinical studies of GBM (Yang et al., 2021). Pre-clinical models of glioma have demonstrated that CD4^+^ T cells are essential for tumor clearance and can induce tumor regression through therapeutic interventions without requiring CD8^+^ T cells (Chen et al., 2022; Kilian et al., 2022; Murphy et al., 2014; Murphy and Griffith, 2016). CD4^+^ T cells can also provide essential help to B cells for effective antibody-mediated immune responses (Eisenbarth et al., 2021; Gutierrez-Melo and Baumjohann, 2023). The presentation of antigens through MHCII is particularly important when CD4^+^ T cell-mediated immune responses are more predominant (Alspach et al., 2019). In line with existing literature (Cantoni et al., 2015; Timperi et al., 2022), our current study showed that TREM2 deficiency reduces the number of MHCII^+^ macrophages and CD4^+^ T cell infiltration. This may explain why we did not observe a significant beneficial effect, but unexpectedly the detrimental consequence of TREM2 deficiency in the pre-clinical model of glioblastoma. In the CD4^+^ T cell dominant context, the benefits of TREM2 deficiency on CD8^+^ T cells may still outweigh the negative effects on CD4^+^ T cells.

We revealed that TREM2 deficiency impaired the ability of myeloid cells to uptake tumor debris, which is the first step in the anti-tumor response through the phagocytosis-MHCII antigen presentation-CD4^+^ axis (**Figure 5L**). The results are consistent with previous research in the CNS, where TREM2 is known to play a key role in phagocytosis of apoptotic neurons (Atagi et al., 2015; Kawabori et al., 2015; Takahashi et al., 2005). In addition, TREM2 function in phagocytosis of protein aggregations has been increasingly recognized in neurodegenerative diseases (Colonna, 2023; Xie et al., 2022b; Zhao and Bu, 2023). Our study is the first to elaborate on the functions of TREM2 in the context of gliomas by demonstrating TREM2-dependent phagocytosis of glioma debris through our *in vivo* imaging and flow cytometry data using mCherry^+^ GL261 and CX3CR1^GFP^ myeloid cells. Based on our findings, we reason that the impaired phagocytosis in TREM2 deficient mice may likely cause the reduced MHCII^+^ macrophages in gliomas. Our findings also provide insight into TREM2 expression beyond the myeloid population, as we identified a modest population of TREM2^+^ T cells in both human GBM and murine GL261. This is especially noteworthy since TREM2 has recently been discovered as a sensor responsible for Th1 activation, and its deficiency in CD4^+^ T cells impairs proinflammatory Th1 responses to infectious diseases (Wu et al., 2021a; Wu et al., 2021b). Therefore, future studies using conditional TREM2 knockout will be required to dissect TREM2 functions in multiple tumor-associated immune cells.

In summary, we have demonstrated that TREM2 play a protective role in gliomas through phagocytosis and antigen presentation. Furthermore, our findings emphasize the importance of evaluating both CD8^+^ and CD4^+^ responses in different tumor contexts when developing TREM2-targeted therapies. As TREM2 antagonists emerge as promising therapeutic targets for cancer treatment, it is crucial to fully understand the range of TREM2 functions in different immune cell types and scrutinize their impact on tumor progression.

## Acknowledgements

The current study is supported provided by the NIH R21AG064159 (LJW), K99 NS1177992 (KA), R01 NS122174 (AJJ), and Mayo Clinic Center for Biomedical Discovery.

---

**Supplementary Figure 1:**
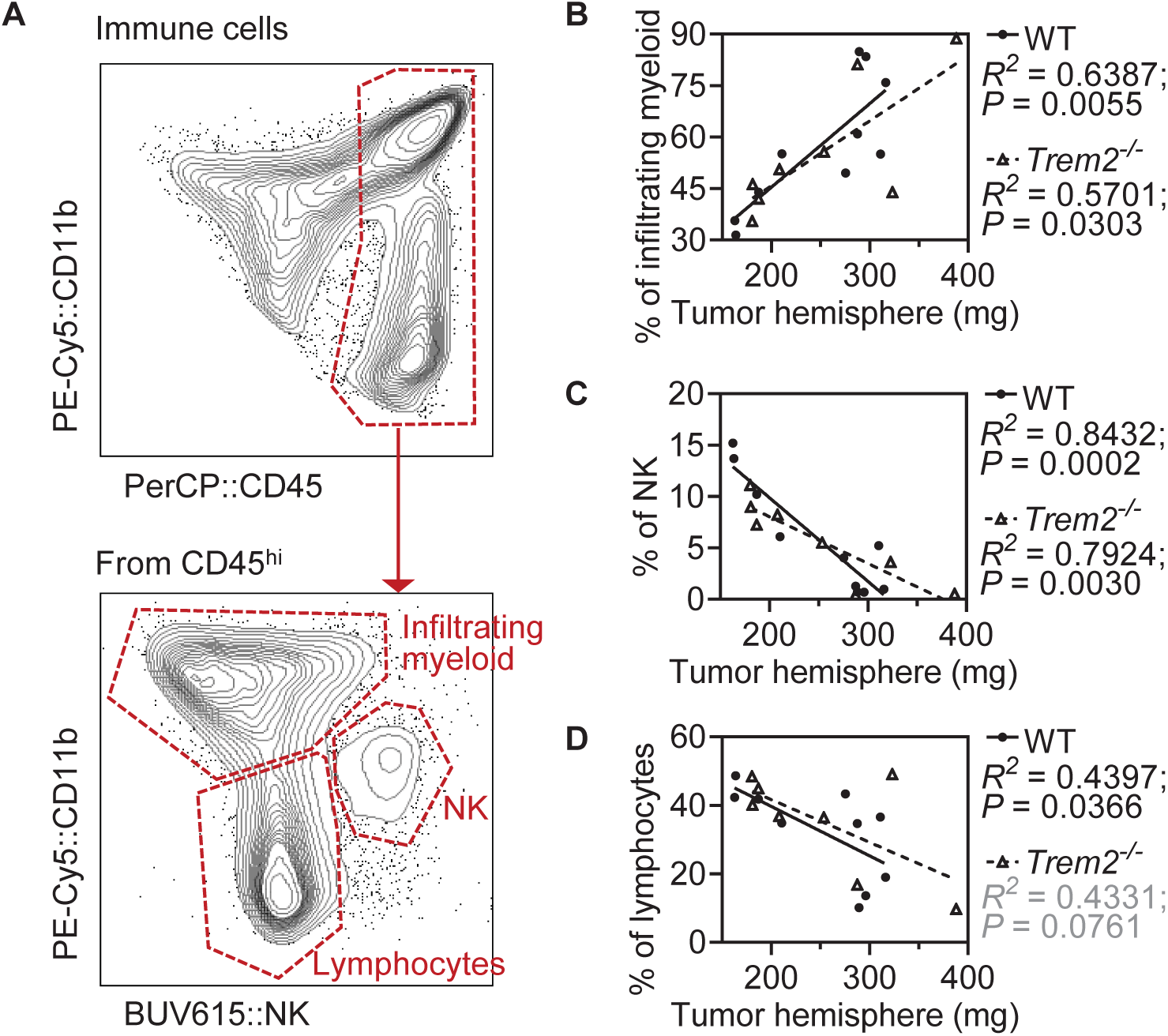
TREM2 deficiency does not alter cellular composition of infiltrating immune cells in brain tumor. **A.** The gating strategy used to identify infiltrating myeloid cells, NK cells, and lymphocytes from CD45^hi^ cluster. **B - D.** Correlation between immune populations and the weight of tumor hemispheres in WT and *Trem2^-/-^*. Spearman’s correlation test was used to calculate *P*-values and correlation coefficients (R^²^).

**Supplementary Table 1:** *TREM2* expression is prevalently elevated in 22 tumor types. A list of 22 tumor types with an increase in *TREM2* expression compared to their paired normal tissues.

| **Tumor types** | **Tumor group sample size** | **Median expression of *TREM2* RNA** | **Non-tumor group sample size** | **Median expression of *TREM2* RNA** |
| --- | --- | --- | --- | --- |
| Glioblastoma (GBM) | 163 | 117.85 | 207 | 2.47 |
| Brain lower grade glioma (LGG) | 518 | 54.32 | 207 | 2.47 |
| Kidney renal clear cell carcinoma (KIRC) | 523 | 26.84 | 100 | 0.92 |
| Kidney renal papillary cell carcinoma (KIRP) | 286 | 26.69 | 60 | 0.82 |
| Pancreatic adenocarcinoma (PAAD) | 179 | 23.80 | 171 | 0.97 |
| Breast cancer (BRCA) | 1085 | 22.63 | 291 | 6.17 |
| Ovarian serous cystadenocarcinoma (OV) | 426 | 12.65 | 88 | 1.27 |
| Head and neck squamous cell carcinoma (HNSC) | 519 | 12.49 | 44 | 1.61 |
| Skin cutaneous melanoma (SKCM) | 461 | 10.98 | 558 | 0.23 |
| Thyroid carcinoma (THCA) | 512 | 10.41 | 337 | 0.97 |
| Cervical squamous cell carcinoma and endocervical adenocarcinoma (CESC) | 306 | 8.80 | 13 | 0.44 |
| Uterine carcinosarcoma (UCS) | 57 | 8.53 | 78 | 0.40 |
| Stomach adenocarcinoma (STAD) | 408 | 8.46 | 211 | 0.44 |
| Testicular germ cell tumors (TGCT) | 137 | 7.00 | 165 | 0.27 |
| Uterine corpus endometrial carcinoma (UCEC) | 174 | 6.98 | 91 | 0.45 |
| Esophageal carcinoma (ESCA) | 182 | 6.02 | 286 | 0.29 |
| Thymoma (THYM), | 118 | 5.84 | 339 | 0.17 |
| Kidney chromophobe (KICH) | 66 | 5.29 | 53 | 0.82 |
| Colon adenocarcinoma (COAD) | 275 | 4.63 | 349 | 0.74 |
| Lymphoid neoplasm diffuse large B-cell lymphoma (DLBC) | 47 | 4.09 | 337 | 0.17 |
| Rectum adenocarcinoma (READ) | 92 | 4.04 | 318 | 0.68 |
| Liver hepatocellular carcinoma (LIHC) | 369 | 3.51 | 160 | 0.25 |

